# Unconscious switching of dorsal and medial pathways for plasticity and stability during NREM and REM sleep

**DOI:** 10.1101/2025.11.04.686485

**Authors:** Takashi Yamada, Theodore LaBonte-Clark, Nitzan Censor, Takeo Watanabe, Yuka Sasaki

## Abstract

While visual perceptual learning improves during non-REM sleep and stabilizes during REM sleep via excitatory-inhibitory neurotransmitter (E/I) balance in early visual areas (EVA), the role of prefrontal regions remains unclear. Here, we show that contributions of the dorsolateral prefrontal cortex (DLPFC) and medial prefrontal cortex (mPFC) differ by sleep stage in human adults. During non-REM sleep, plasticity increased in DLPFC—indexed by elevated E/I balance measured with magnetic resonance spectroscopy and polysomnography—in correlation with performance gains. During REM sleep, stability increased in mPFC—indexed by reduced E/I balance—in correlation with resilience to retrograde interference from new learning. E/I balance changes and their effects on learning paralleled those in EVA. Connectivity weights between EVA and DLPFC, and between EVA and mPFC, switched with sleep stage. These findings suggest the presence of dorsal and medial pathways that unconsciously alternate between non-REM sleep and REM sleep to improve and stabilize learning.

## INTRODUCTION

Sleep facilitates learning and memory (*1–8*), collectively referred to as *sleep-facilitated effects*. These effects can be divided into two complementary types. First, sleep enhances performance after sleep compared with before, a phenomenon known as offline performance gains (*2, 4, 5, 8*). Second, sleep stabilizes previously acquired learning by protecting it from retrograde interference (*9, 10*), in which new learning disrupts earlier knowledge (*11, 12*).

To uncover the neural mechanisms underlying these sleep-facilitated effects, it is crucial to recognize that sleep comprises two distinct stages—non-rapid eye movement (NREM) sleep and rapid eye movement (REM) sleep—each characterized by unique patterns of neural activity across cortical and subcortical regions (*13–17*). Once considered a state of neural quiescence, sleep is now known to involve dynamic and sleep-stage specific reorganization of brain activity. PET and fMRI studies revealed that regional activation can increase during both NREM and REM sleep, indicating that sleep is not a passive but an active process that reorganizes functional networks to support learning and memory.

Among brain regions implicated in these processes, frontal, control-related areas such as the prefrontal cortex (PFC) are thought to play central roles. The dorsolateral PFC (DLPFC) shows increased activation during NREM sleep following visual training (*8*), whereas the medial PFC (mPFC) becomes more active during REM Sleep (*13, 17, 18*). However, how these regions contribute differentially to sleep-facilitated effects — and how they coordinate with other regions including sensory areas—remains unsolved.

Inspired by a series of work on developmental physiology (*19*), which demonstrated that the excitatory–inhibitory (E/I) balance governs the onset and offset of the visual critical period, subsequent research has shown that plasticity and stability in post-training stages of learning and memory are closely associated with the E/I. In human studies, the concentration of Glx (a combined signal of glutamate, an excitatory neurotransmitter, and glutamine) and GABA, an inhibitory neurotransmitter have been extensively measured by magnetic resonance spectroscopy (MRS) (*20–22*). Recently, a number of studies have calculated Glx divided by GABA as the E/I ratio (*23, 24*). This relationship is particularly robust when the E/I ratio is assessed at multiple time points within the same individual—an approach known as the functional E/I ratio (fE/I ratio). Such a within-subject design minimizes interindividual variability and captures dynamic, state-dependent changes in the E/I ratio. Notably, increases in the fE/I ratio are highly correlated with plasticity, as reflected by performance enhancement, whereas decreases in the fE/I ratio are associated with stability, as reflected by resilience to retrograde interference (*5, 24–31*).

In this study, we investigated how the DLPFC and mPFC differently contribute to offline performance gains (plasticity) and stabilization (stability) across sleep stages. Using simultaneous MRS and polysomnography, we measured Glx and GABA during NREM and REM sleep relative to wakefulness using and calculated fE/I ratios to examine how these neurochemical dynamics relate to learning outcomes. We further explored functional links between prefrontal subregions and early visual areas, which likely represent plasticity in visual learning, to delineate large-scale pathways underlying offline performance gains and stabilization.

We found that the DLPFC and mPFC play distinct yet complementary roles. Following orientation detection training, the DLPFC showed increased E/I ratios during NREM sleep, which correlated with offline performance gains, whereas the mPFC showed decreased E/I ratios during REM sleep, which correlated with stabilization. Notably, these bidirectional changes mirrored patterns previously observed in early visual areas (*5*). Together with analyses of gamma-connectivity—gamma band activity being closely linked to E/I ratios—, these results suggest the existence of two functionally and anatomically district pathways: one pathway, active during NREM sleep involving the DLPFC and early visual areas, increases E/I ratio, and supports offline performance gains. The other pathway, active during REM sleep involving the mPFC and early visual areas, deceases the E/I ratio and contributes to sleep stabilization. Since NREM and REM cycles alternate automatically throughout human sleep, independent of conscious involvement (*31*), the dominance of these two pathways may switch across cycles, operating without conscious control.

## RESULTS

To investigate the roles of the DLPFC and mPFC in offline performance gains and stabilization (resilience to retrograde interference), we conducted two distinct experiments. In the first experiment, we examined the DLPFC, whereas in the second experiment, we focused on the mPFC. Using EEG data from both experiments, we also calculated gamma-band connectivity between the PFC (DLPFC and mPFC) and early visual areas.

### Experiment 1

The role of the DLPFC in offline performance gains and stabilization during NREM and REM sleep was investigated in 15 participants (10 females).

As shown in Figure 1A, the complete experiment consisted of two weeks of sleep monitoring, one adaptation stage, and one day of training and measurement. During the sleep monitoring stage, each participant’s sleep quality was assessed using sleep logs and/or actigraph devices to record daily sleep patterns. Based on these measurements, the timing of the training and measurement stage was scheduled to ensure each participant experienced normal sleep (see Experimental design in the methods).

**Figure 1.**
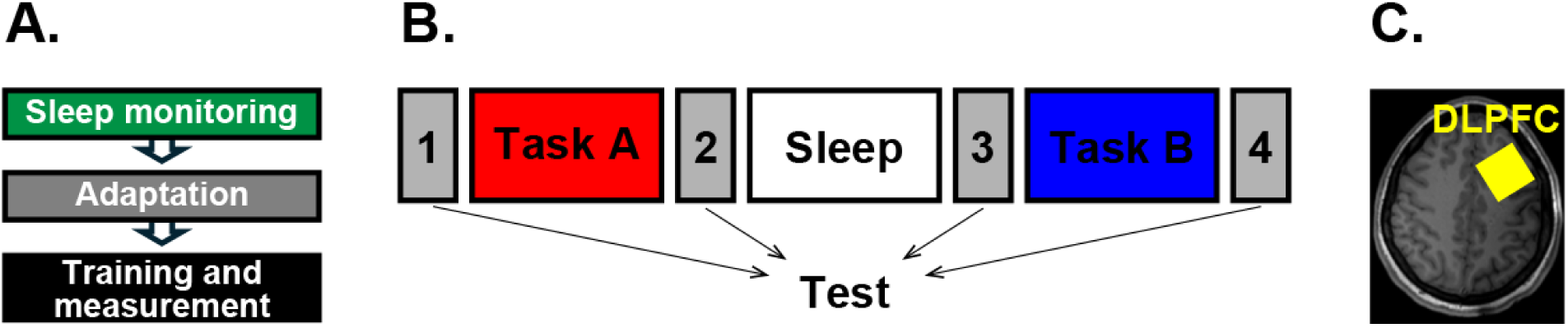
Procedures of Experiment 1. (A) Stages of the complete experiment. (B) Procedures of the training sessions (Tasks A and B) and test stages. C. Voxels of interest (VOIs) placed in the DLPFC.

During the adaptation stage, participants spent one night in the sleep laboratory to experience the first night effect (FNE), thereby minimizing its influence on post-training sleep. The FNE (*32–37*) is characterized by significantly lower sleep quality, particularly during the first night in a new environment. After an interval of at least one week following the adaptation stage, participants completed the training and measurement stage, which lasted one day. During this stage, we investigated whether and how the DLPFC is involved in offline performance gains and resilience to retrograde interference.

To obtain the degree of offline performance gains, we measured how performance on the trained task changed from before to after sleep that followed visual training. First, the Training A stage occurred. In this stage, participants were asked to perform the texture discrimination task (TDT), which is a highly standard test used for VPL studies. In each trial, after participants performed the first task to ensure their gaze at the center of the display (see Texture discrimination task (TDT) in the Methods), they were asked to report whether a target consisting of three segments whose orientation differed from that of the rest of the segments in the display was vertically or horizontally elongated. The target was consistently presented in the upper left quadrant of the display or in the upper right quadrant, randomly assigned to each participant. The target was followed by a blank display and then a mask disrupting the performance on the target. The stimulus-to-mask onset asynchrony (SOA) that gives the 80% target performance was determined as the threshold performance. Test 2 and Test 3 were conducted before and after sleep, respectively, following the same procedure. They consisted of 90 trials of the same TDT task as in Training A. The degree of performance gains (%) was defined by (threshold in Test 2-threshold in Test 3)/ threshold in Test 1 x 100.

To measure the degree of resilience to retrograde interference, Training B and Test 4 were further conducted. The procedures of Training B were the same as Training A, with the following exception: the orientation of the background elements was orthogonal to that in the stimulus of Training A. This setting was made because a previous study found that retrograde interference from VPL of Training B to VPL of Training A occurred when Training A used the TDT stimuli whose background elements’ orientations were orthogonal to those in the TDT stimuli Training B used (*38*). After Training B, Test 4 was conducted with the same procedure as Tests 2 and 3. The degree of resilience to retrograde interference (%) was defined by (threshold in Test 4-threshold in Test 3)/threshold in Test 3 x 100.

Figure 2 shows the mean (±s.e.m.) degree of offline performance gains (left) and resilience to retrograde interference (right). Since data measurements such as E/I ratios and gamma connectivity in some conditions violated normality (see below), we used the bootstrapping analysis throughout the data of this study to unify the analysis method. *Here, the* analysis, based on 2,000 resamples, yielded the following results: both offline performance gains was greater than zero (*t*=4.86, *p*<0.001, 95% CI=[7.81, 18.01], Cohen’s *d*=1.25). In contrast, resilience to retrograde interference did not reach statistical significance (*t*=1.56, *p*=0.12, 95% CI=[-1.41, 12.59], Cohen’s *d*=0.40), which indicates performance did not get worse despite the competing task training. These results show that sleep contributed to both the degree of offline performance gains and resilience of retrograde interference.

**Figure 2.**
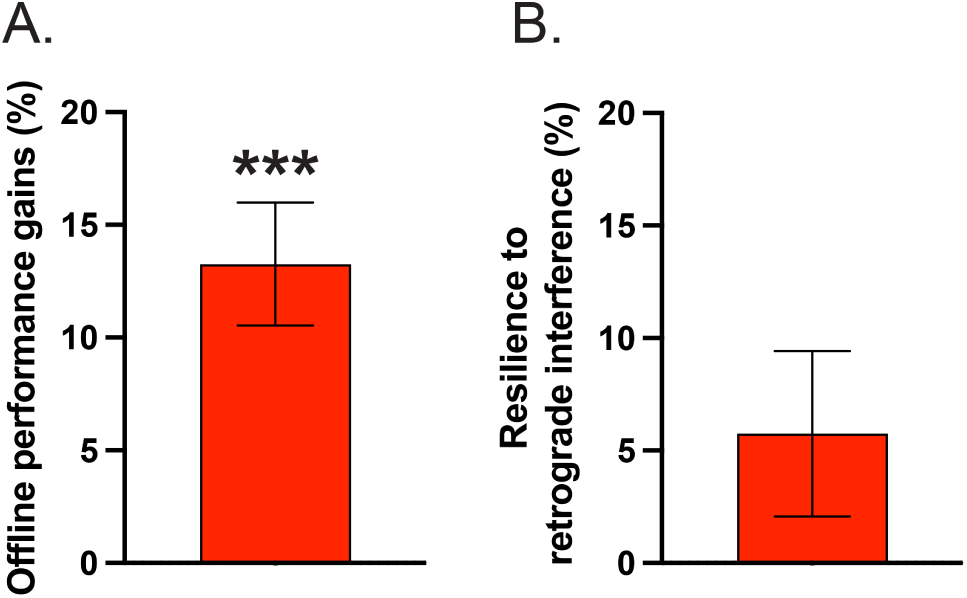
Behavioral results of Experiment 1. A. Mean (±s.e.m.) offline performance gains (%), defined as [(Performance in Test 2-Performance in Test 1)/(Performance in Test1)] x 100. B. Mean (±s.e.m.) resilience to retrograde interference (%), defined as [(Performance in Test 3 -Performance in Test 2)/(Performance in Test 2)] x 100.

During the sleep stage, participants were asked to lay down in a magnet booth and sleep. Before and after participants fell asleep in the magnet, the concentrations of Glx (glutamate + glutamine) and GABA in the DLPFC were measured using magnetic resonance spectroscopy (MRS) simultaneously with polysomnography measurements that determined sleep stages.

The E/I ratio in the respective VOI was calculated during wakefulness prior to sleep, NREM sleep and REM sleep. Each E/I ratio was defined by the concentration of Glx divided by the concentration of GABA. By calculating the change of E/I ratio during non-REM or REM sleep relative to E/I ratio during wakefulness, we obtained fE/I ratios, which represents changes in the E/I ratio relative to the baseline wakefulness stage to NREM sleep or REM sleep.

Figures 3 show the mean (±s.e.m.) E/I ratio change from wakefulness to NREM sleep and REM sleep for the DLPFC. The Shapiro–Wilk tests showed that the data for NREM sleep met the normality assumption (W=0.91, p=0.15) but not for REM sleep (W=0.80, p=0.03). To unify the method, we conducted the bootstrap analysis. Based on 2,000 resamples, the results yielded the following results: the mean E/I ratio change from wakefulness to NREM sleep was significant (*p*=0.005, Cohen’s *d*=0.66, 95% CI=[2.30, 15.06]), whereas the change from wakefulness to REM sleep was not significant (*p*=0.81, Cohen’s *d*=-0.04, 95% CI=[-9.05, 10.50]).

**Figure 3.**
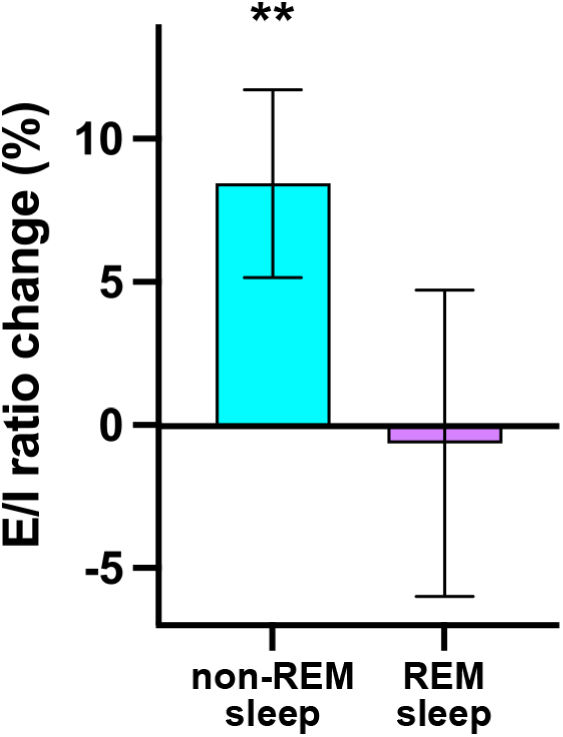
Mean (±s.e.m.) E/I ratio changes from wakefulness to NREM sleep (left) and REM sleep (right) in DLPFC.

We further investigated how the E/I ratio in the DLPFC is associated with offline performance gains and resilience to retrograde interference. Given that the E/I ratio change in the DLPFC was significant only from wakefulness to NREM sleep, we examined the correlation between this E/I ratio change and offline performance gains and that between the E/I ratio changes and resilience to retrograde interference across participants. As shown in Figure 4A and 4B, the bootstrap analysis with 2,000 iterations shows that the E/I ratio change from wakefulness to NREM sleep significantly correlated with offline performance gains (*r*=0.563, *t*=2.456, *p*=0.002, 95% CI=[0.236, 0.818]) but not with resilience to retrograde interference (*r*=-0.030, *t*=-0.108, *p*=0.958, 95% CI=[-0.649, 0.539]). Additionally, as shown in Figure 4C and 4D, the E/I ratio change from wakefulness to REM sleep did not significantly correlate with either offline performance gains (*r*=-0.171, *t*=-0.424, *p*=0.493, 95% CI=[-0.866, 0.491]) or resilience to retrograde interference (*r*=0.510, *t*=1.453, *p*=0.579, 95% CI=[-0.561, 0.978]).

**Figure 4.**
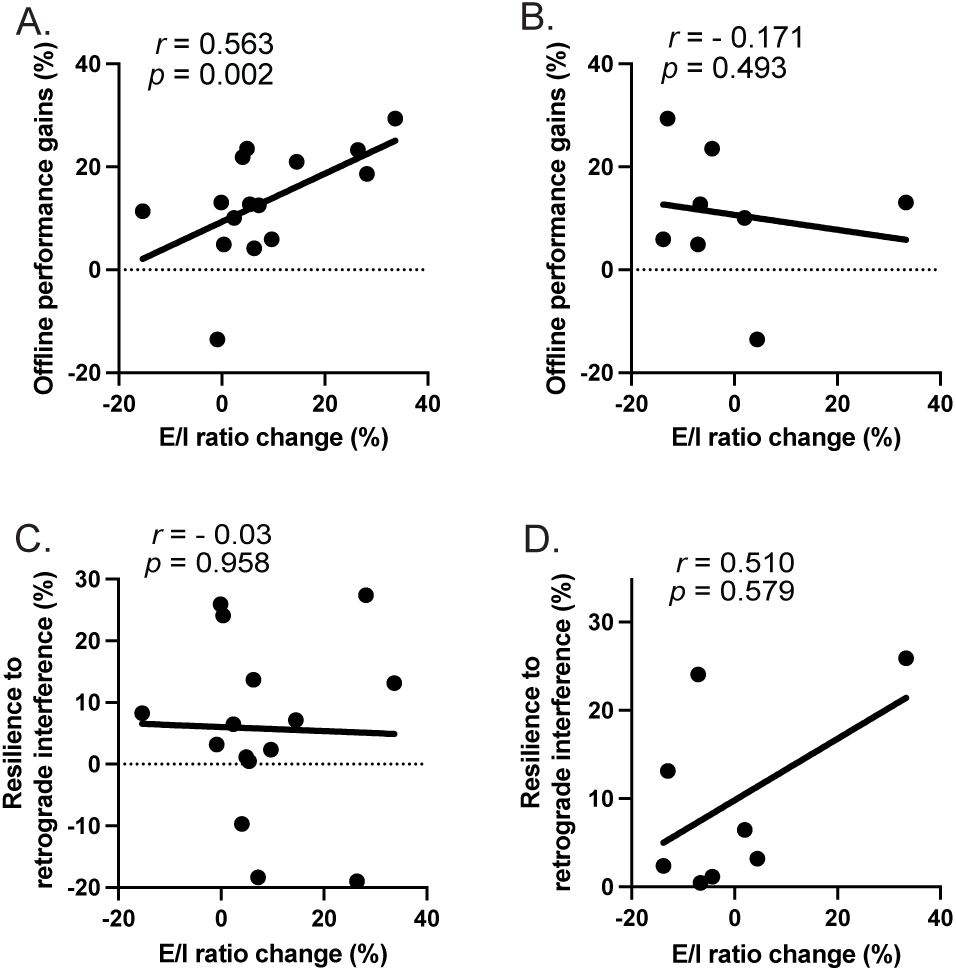
Scatter plots in Experiment 1. A. Offline performance gains vs. changes in E/I ratio in DLPFC from wakefulness to NREM sleep. B. Offline performance gains vs. changes in E/I ratio in DLPFC from wakefulness to REM sleep. C. Resilience to retrograde interference gains vs. changes in E/I ratio in DLPFC from wakefulness to NREM sleep. D. Resilience to retrograde interference gains vs. changes in E/I ratio in DLPFC from wakefulness to REM sleep. Solid lines represent linear regression using the least squared method.

### Experiment 2

In this experiment, a new group of 16 participants (9Females) were recruited to measure the concentrations of Glx and GABA in the mPFC. Two separate experiments (Experiments 1 and 2) were conducted to measure the concentrations in different regions, as our experimental procedures do not allow for simultaneous measurement of two regions.

Figure 5 shows the mean (±s.e.m.) degree of offline performance gains (left) and resilience of retrograde interference (right). We applied the bootstrapping analysis, in order to unify the method with Experiment 1. Based on 2,000 resamples, this analysis yielded the following results: offline performance gains were significantly greater than zero (*t*=4.67, *p*<0.001, 95% CI=[5.50, 13.00], Cohen’s *d*=1.17), whereas resilience to retrograde interference did not reach significance (*t*=1.59, *p*=0.11, 95% CI=[-1.25, 13.22, Cohen’s *d*=0.41). These results indicate that sleep contributed to offline performance gains and resilience of retrograde interference, consistent with the tendencies observed in Experiment 1.

**Figure 5.**
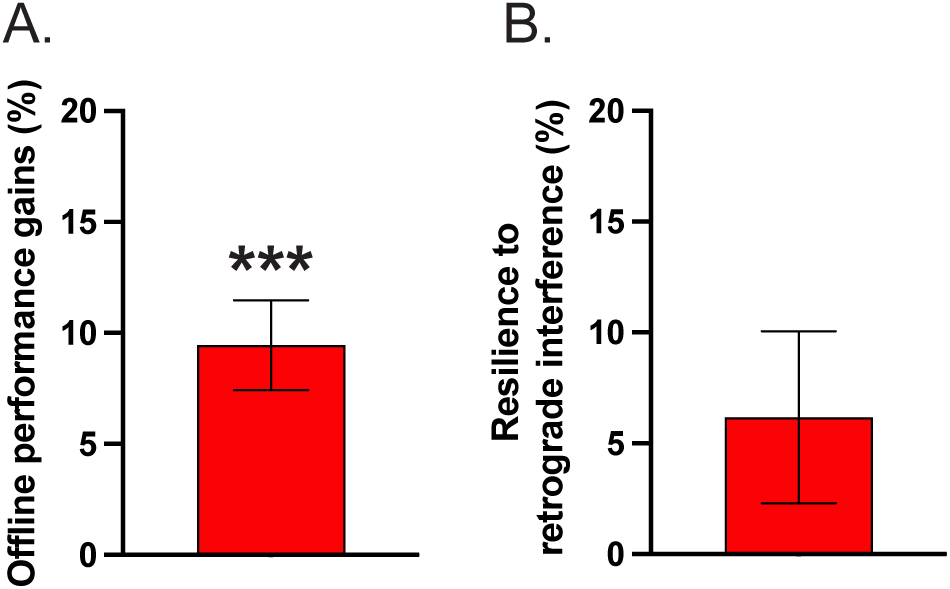
Behavioral results of Experiment 2. A. Mean (±s.e.m.) offline performance gains (%), defined as [(Performance in Test 2-Performance in Test 1)/(Performance in Test1)] x 100. B. Mean (±s.e.m.) resilience to retrograde interference (%), defined as [(Performance in Test 3 -Performance in Test 2)/(Performance in Test 2)] x 100.

As mentioned above, the concentrations of Glx and GABA were measured in the mPFC (Figure 6A). Although the data did not violate the normality assumption [W=0.93. p=0.23 for NREM; W=0.89, p=0.13 for REM], we conducted bootstrapping analysis to unify the method with Experiment 1 where the data violated the normality assumption. In contrast to the DLPFC, the results indicated that E/I ratio change in the mPFC, from the wakefulness to NREM sleep was not significant (*p*=0.272, Cohen’s *d* = 0.26, 95% CI = [-3.39, 12.31]), whereas that from wakefulness to REM sleep was significant (*p*=0.01, Cohen’s *d* = -0.70, 95% CI = [-18.90, -2.84]).

**Figure 6.**
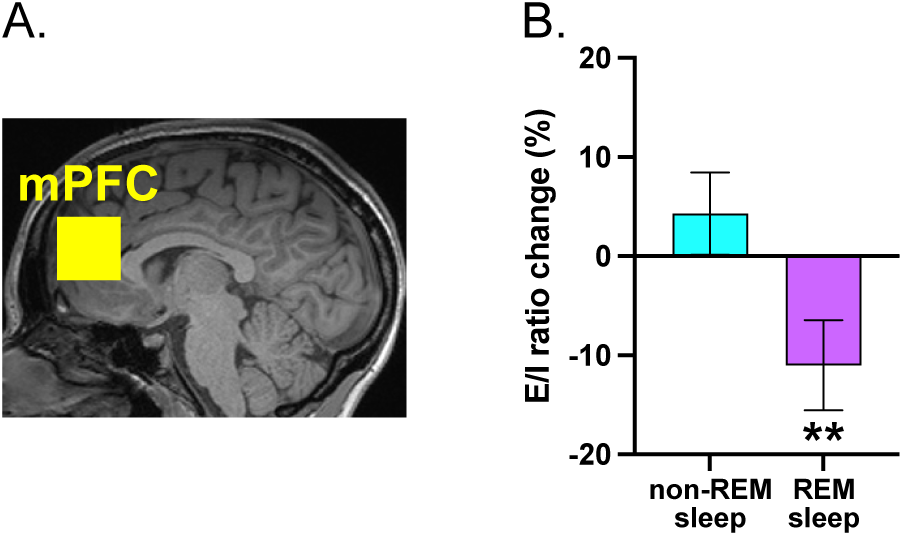
A. VOI placed in mPFC. B. Mean (±s.e.m.) E/I ratio changes from wakefulness to NREM sleep (left) and REM sleep (right) in mPFC.

Associations of the E/I ratio in the mPFC with offline performance gains and resilience to retrograde interference showed the opposite tendency to those in the DLPFC. As shown in Figure 7A and B, bootstrap analysis with 2,000 iterations indicates that the E/I ratio change from wakefulness to NREM sleep did not significantly correlate either with offline performance gains (*r*=0.283, *t*=1.103, *p*=0.291, 95% CI=[-0.264, 0.685]) or resilience to retrograde interference (*r*=-0.261, *t*=-0.975, *p*=0.319, 95% CI=[-0.852, 0.281]). Conversely, the E/I ratio change from wakefulness to REM sleep significantly inversely correlated with resilience to retrograde interference (*r*=-0.654, *t*=-2.737, *p*=0.003, 95% CI=[-0.891, -0.279]) but not with offline performance gains (*r*=-0.311, *t*=-1.034, *p*=0.207, 95% CI=[-0.729, 0.217]).

**Figure 7.**
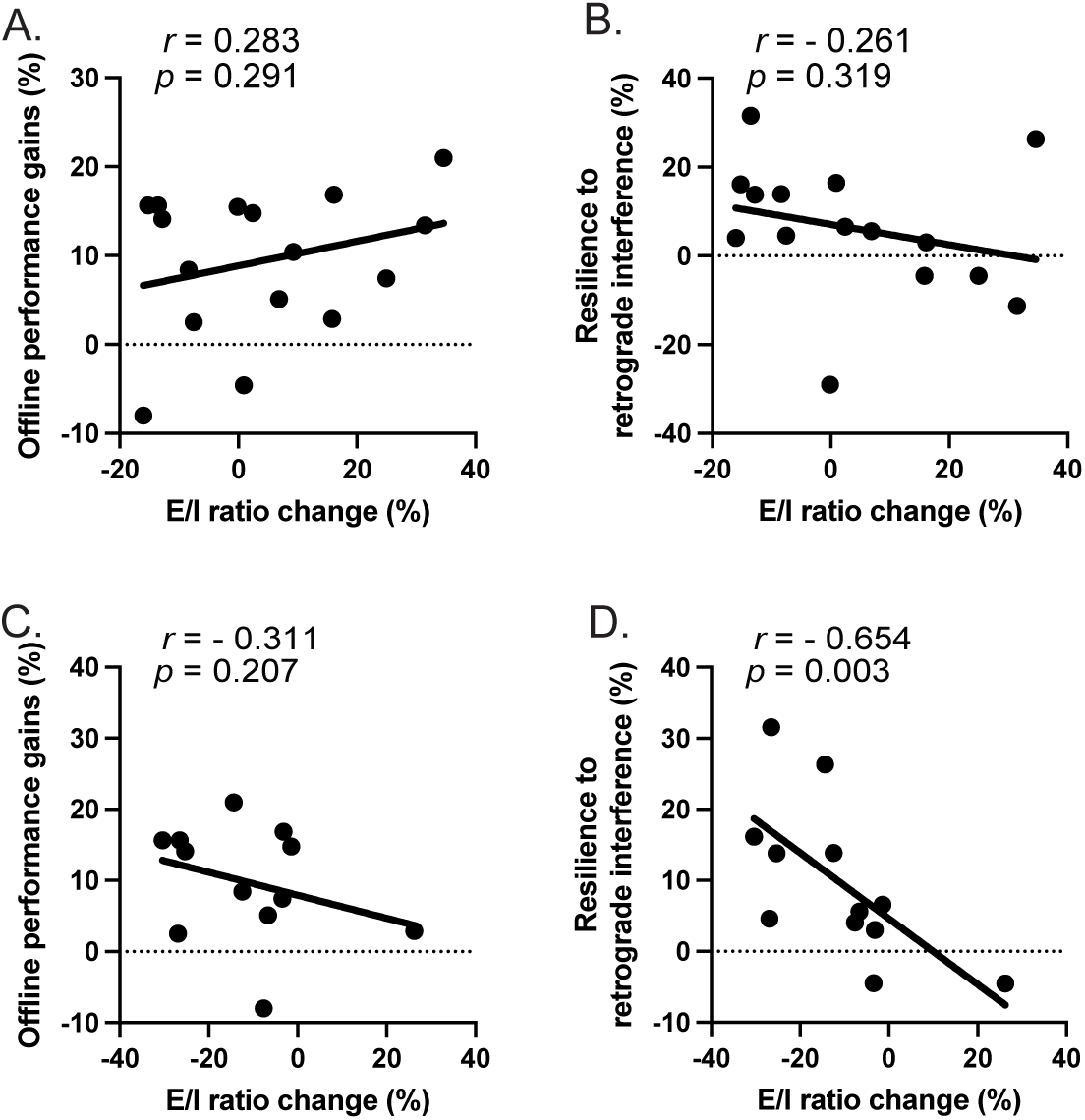
Scatter plots in Experiment 2. A. Offline performance gains vs. changes in E/I ratio in mPFC from wakefulness to NREM sleep. B. Resilience to retrograde interference gains vs. changes in E/I ratio in mPFC from wakefulness to NREM sleep. C. Offline performance gains vs. changes in E/I ratio in mPFC from wakefulness to REM sleep. D. Resilience to retrograde interference gains vs. changes in E/I ratio in mPFC from wakefulness to REM sleep. Solid lines represent linear regression using the least squared method.

These results suggest the different roles of DLPFC and mPFC in learning sleep during different sleep stages. During NREM sleep, DLPFC appears to be predominantly involved in performance gains associated with an increased E/I ratio, whereas during REM sleep, mPFC may plays a role in sleep stabilization associated with a decreased E/I ratio.

### Two pathways

Interestingly, our previous study measured the E/I ratio in early visual areas under conditions involving sleep stages and visual learning mirror those in the PFC in the current study (*5*). The E/I ratio in early visual areas significantly increased during NREM sleep and was strongly correlated with the degree of offline performance gains. In contrast, the E/I ratio significantly decreased during REM sleep and was inversely correlated with resilience to retrograde interference. These results suggest a similar role of the DLPFC and early visual areas in facilitating offline performance gains during NREM sleep, and a comparable role of the mPFC and early visual areas in enhancing resilience to retrograde interference during REM sleep.

This raises the possibility that during NREM sleep, the DLPFC and EVA coordinate to support offline performance gains, whereas the mPFC and EVA work together for sleep stabilization. Among commonly used oscillatory measures, gamma-band power appears to best reflects the E/I ratio (*39, 40*). Thus, we investigated whether DLPFC-EVA gamma connectivity and mPFC-EVA gamma connectivity exhibit different patterns during NREM and REM sleep. We used Low-Resolution Electromagnetic Tomography (LORETA) to estimate source-level activity from EEG signals recorded during NREM and REM sleep in Experiments 1 and 2, and we calculated gamma-band connectivity between EVA and prefrontal regions, such the DLPFC and mPFC. Compared with analyzing raw gamma activity, applying LORETA significantly increases the reliability of estimating cortical gamma-band activity and connectivity. While EEG is less suited for detecting local or deep structures, we took the 30-40hz range which is much less noisy than higher frequencies and our focus on global cortical coupling ensures our method an informative approach (*41, 42*).

The results are shown in Fig. 8. In contrast to the MRS data in which only one region was measured for each participant, gamma connectivity was assessed in both DLPFC-EVA and mPFC-EVA pathways for all participants. The Shapiro–Wilk test indicated none of the d from normality. In contrast, the normality assumption was rejected for mPFC-EVA connectivity during REM sleep (W=0.91, p=0.035). We applied a 2-way repeated-measures ANOVA with bootstrap resampling (2,000 iterations) to gamma connectivity, with Pathway (DLPFC-EVA vs. mPFC-EVA) and Sleep Stage (NREM vs. REM) as within-subject factors. The analysis revealed a significant interaction between the two factors [*F*=26.29, *p*<0.001, partial *η*^2^=0.56]. Given this significant interaction, we conducted post-hoc paired-sample t tests with bootstrap resampling (2,000 iterations). The test showed that DLPFC-EVA gamma connectivity was significantly lower during NREM sleep than REM sleep [*t*=-2.96, *p*=0.0085, Cohen’s *d*=-0.63]. Conversely, the results of the mPFC-EVA pathway showed that gamma connectivity was significantly higher during NREM sleep than REM sleep [*t*=3.05, *p*=0.0055, Cohen’s *d*=0.65]. These results suggest the involvement of two distinct pathways: a dorsal pathway, including the DLPFC and EVA, which is activated during NREM sleep and associated with offline performance gains, and a medial pathway, involving the mPFC and EVA, which is activated during REM sleep with resilience and associated with retrograde interference (see Fig.9).

**Figure 8.**
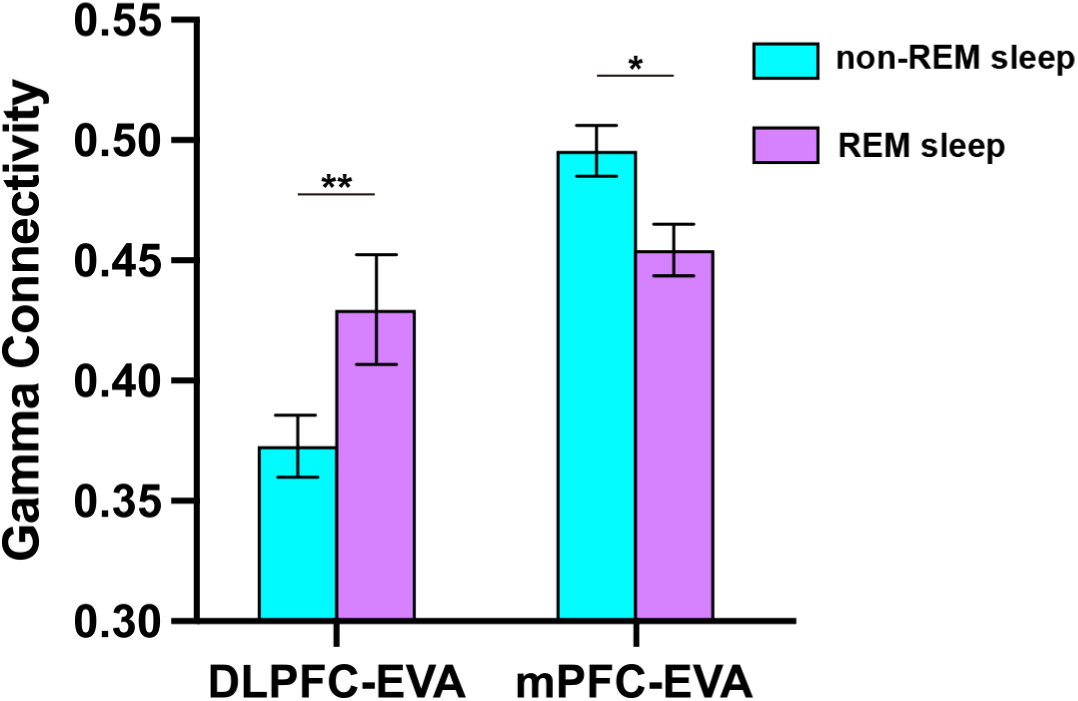
Mean (±s.e.m.) Gamma connectivity between DLPFC and EVA, and between mPFC and EVA, during REM and non-REM sleep. Asterisks indicate significant differences (** P<0.01, * P<0.05).

**Figure 9.**
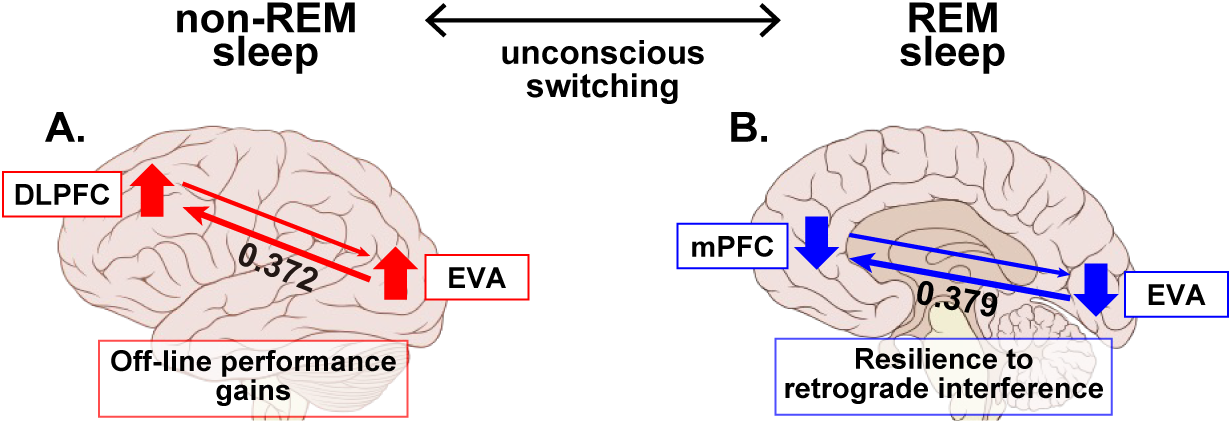
Schematic illustration of two pathways between the prefrontal and early visual areas (EVA) with corresponding Granger causality (GC) values. A. Pathway between the DLPFC and EVA, supporting offline performance gains during NREM sleep, which are associated with an increased E/I ratio. GC values from EVA to DLPFC and from DLPFC to EVA are shown beside each arrow. B. Pathway between the mPFC and EVA, contributing to stabilization (resilience to retrograde interference) during the REM sleep, which are associated with a decreased E/I ratio. GC values from EVA to mPFC and from mPFC to EVA are shown beside each arrow.

To infer the causal flows of signal processing between the DLPFC and EVA during NREM sleep, as well as between the mPFC and EVA during NREM sleep, we conducted Granger causality (GC) analysis (see PSG measurement). Only 23 of the 30 subjects were included in the analysis because they exhibited both NREM and REM sleep. <u>A</u> one-way repeated-measures ANOVA with bootstrap resampling (2,000 iterations) revealed a significantly larger GC from EVA to DLPFC (0.151) than from DLPFC to EVA (0.112) during NREM sleep (*F*=9.229, *p*=0.011, Cohen’s *d*=-0.550, 95% CI=[0.064, 0.015]). These results suggest stronger bottom-up signal propagation from EVA to the DLPFC during NREM sleep.

In contrast, during REM sleep, GC from EVA to mPFC (0.153) was significantly larger than from mPFC to EVA (0.131) (*F*=9.295, *p*=0.007, Cohen’s *d*=-0.636, 95% CI=[0.066, 0.017]), indicating enhanced bottom-up communication from EVA to the mPFC. GCs of both EVA↔DLPFC during NREM sleep and EVA↔mPFC during REM sleep fell within the moderate range typical of inter-areal EEG connectivity (10–20% improvement in predictability).

Overall, these findings demonstrate that signal proceeding between EVA and each prefrontal region is bidirectional. However, the slightly but significantly higher GC values in the bottom-up direction indicate that communication from EVA to the PFC is more dominant than in the reversed direction, suggesting that visual cortical activity may exert a stronger influence on prefrontal regions during both NREM and REM sleep.

## GENERAL DISCUSSION

We investigated whether and how two prefrontal regions—the DLPFC and mPFC— facilitate learning by examining changes in E/I ratios in these regions from wakefulness to NREM and REM sleep. The results showed that the DLPFC plays a significant role in offline performance gains, which are associated with increased E/I ratios from wakefulness to NREM sleep, whereas the mPFC is primarily involved in stabilization indexed by the resilience to retrograde interference, which is associated with decreased E/I ratios from wakefulness to REM sleep.

Interestingly, a study found that the direction of E/I ratio changes and their associated sleep stages mirrored those observed in EVA (*5*). Therefore, we examined gamma connectivity between the DLPFC and EVA, as well as between and the mPFC and EVA. We found distinct patterns of connectivity between NREM and REM sleep for the DLPFC-EVA and mPFC-EVA pathways. Collectively, these results suggest that the DLPFC-EVA pathway supports performance gains during NREM sleep, whereas the mPFC-EVA pathway supports stabilization during REM sleep. In humans, NREM and REM sleep cycles alternate repeatedly without conscious awareness (*43, 44*), this cycling may reflect a dynamic shift in dominance between these two pathways across sleep stages, functioning unconsciously.

DLPFC–EVA and mPFC–EVA gamma connectivity exhibited distinct patterns across sleep stages, consistent with the different directions of E/I ratio changes—positive in both the DLPFC and EVA during NREM sleep and negative in both the mPFC and EVA during REM sleep. A question arises: why was DLPFC–EVA gamma connectivity lower during NREM than REM, whereas mPFC–EVA gamma connectivity was lower during REM than NREM?

A likely explanation lies in the nonlinear relationship between E/I balance and gamma activity. Moderate increases in excitatory drive enhance gamma power, but excessive excitation desynchronizes firing and diminishes gamma amplitude (*39*). Likewise, moderate inhibitory drive via PV+ interneurons improves pyramidal timing and strengthens gamma power, whereas excessive inhibition suppresses activity and weakens oscillations (*45*). When local gamma power decreases under either extreme, interregional phase-locking becomes unreliable, reducing long-range synchronization. During NREM sleep, moderately dominant excitation may promote synchronized gamma activity between DLPFC and V1, yielding positive E/I ratios in both; during REM sleep, moderately dominant inhibition may enhance mPFC–V1 synchrony, leading to negative E/I ratios. However, when excitation or inhibition becomes excessive, gamma power and coupling both decline—explaining why DLPFC–V1 connectivity is weaker in NREM and mPFC–V1 connectivity weaker in REM. However, Further research is needed to directly test this possibility.

Recent work (*46*) challenged the assumption that MRS-based E/I ratios universally reflect a stable balance between excitatory and inhibitory neurotransmission. They reported no significant correlation between Glx and GABA concentrations across individuals in either visual or motor cortices, suggesting that trait-level coupling between these metabolites is not a general property of the human cortex. However, as discussed above, the present study does not rely on cross-sectional coupling but instead examines functional E/I (fE/I) ratios—within-subject changes in the Glx/GABA ratio measured across different brain states (e.g., wakefulness, NREM sleep, and REM sleep). Because each individual serves as their own baseline, this approach mitigates confounds arising from between-subject variability in voxel composition, baseline metabolite pools, and measurement noise. Thus, the fE/I ratio used here provides a state-dependent index of dynamic shifts in plasticity and stability, circumventing the major criticism raised by the study (*46*) that static E/I ratios lack general validity across individuals.

Although MRS captures metabolite signals from both intra- and extracellular environments, the observed variations in Glx and GABA levels in our study likely represent, at least in part, changes in synaptic transmission. Previous research has demonstrated that manipulations known to influence glutamatergic or GABAergic synaptic activity—such as pharmacological interventions (*47*), transcranial direct current stimulation (*20, 48, 49*), and engagement in memory-related tasks (*50, 51*)—lead to measurable alterations in MRS-derived Glx and GABA concentrations. Moreover, both perceptual learning (*5, 25, 26, 30, 31*) and sleep (*5, 52–54*) have been shown to modulate excitatory and inhibitory neural activity. Therefore, it is reasonable to interpret the Glx and GABA changes observed in our experiment as indicators of synaptic transmission dynamics, given that these differences emerged in EVA and PFC across distinct sleep stages following visual training, while other factors were kept constant within individuals.

In declarative memory, interactions between the hippocampus and the cortex appear to play significant roles in both offline performance gains and resilience to retrograde interference. Extensive research suggests that, during sleep, the hippocampus reactivates memory traces and interacts with the cortex, strengthening memories as offline performance gains and enhancing resilience to retrograde interference (*55, 56*). However, it remains unclear whether and how these distinct functions—offline performance gains and resilience to retrograde interference —are processed differently. This unclarity may, at least partially, arise from complex interactions among the hippocampus, PFC, and cortical areas where memory is transferred from the hippocampus for long-term storage. Unlike episodic memory, the hippocampus does not appear to play a significant role in VPL. This distinction supports the viability of a simpler, two-pathway processing model for sleep facilitation in VPL.

The present study found that two specific regions in the prefrontal cortex—the DLPFC and mPFC—play distinct roles in sleep-facilitated learning, supporting offline performance gains and stabilization against retrograde interference, respectively. This functional dissociation may reflect their distinct yet interactive contributions to the plasticity–stability trade-off in learning. The DLPFC, known for top-down control and goal-directed modulation of sensory processing (*57–59*), may promote offline performance gains during NREM sleep by enhancing excitatory drive and coordinating reactivation of task-relevant visual representations (*5, 60, 61*). Importantly, this pathway may also involve bottom-up inputs from early visual areas, providing error- or salience-related sensory signals that guide DLPFC-driven reorganization of cortical representations. In contrast, the mPFC, implicated in valuation and integration (*55, 62*), may support stabilization during REM sleep through inhibitory dominance that suppresses competing traces and consolidates strengthened ones (*5, 63, 64*). Our results further suggest that both top-down and bottom-up interactions operate within these pathways, with slightly stronger bottom-up influence. Such reciprocal communication indicates that sleep-stage-specific learning optimization arises from dynamic bidirectional coordination between prefrontal and visual cortices, implementing the plasticity–stability balance fundamental to memory consolidation (*65, 66*).

## METHODS

### Participants

Following careful screening (see below), 31 young and healthy adults (19 females, aged 18 to 30) participated in this study. All the participants provided their written informed consent after being fully briefed on the study’s objectives, procedures, and any potential risks involved, including exposure to loud acoustic noise, possible warmth from radiofrequency energy, and discomfort in confined spaces related to MRI scans. The Institutional Review Board at Brown University approved the study protocol.

To estimate the number of participants required to reliably indicate the effect of a nap on the correlations between the offline performance gains of VPL and E/I ratio and between the degree of resilience to retrograde interference and E/I ratio, we applied the G*Power software (*67, 68*)—set to a power of 0.8, a required significance level (α) of 0.05, and two-tailed—to our published data (*5*). The result indicated 14 as the minimum sample size for the correlation between offline performance gains and E/I ratio and 10 as the minimum sample size for the correlation between the degree of resilience to retrograde interference and E/I ratio. Consequently, we recruited 15 -16 participants per experiment, each focusing on one prefrontal region (either DLFPC or mPFC). This strategy accounted for the potential involvement of both regions in offline performance gains and ensured at least 14 data points in both prefrontal region experiments, even after excluding ineligible participants or outliers.

### Screening

We carefully screened potential participants using established methods to ensure eligibility for both VPL and sleep experiments (*5, 37, 69–71*). Our eligibility criteria focused on the following five key areas: (i) Vision. Participants were required to have normal or corrected-to-normal vision. (ii) VPL. Participants were required to have no prior experience with VPL tasks. This requirement was crucial, as previous exposure can lead to long-term changes in visual sensitivity (*72–78*). (iii) Video Game Experience. We excluded frequent action video game players (defined as those playing more than 5 hours per week for more than 6 months) due to the potential effects on visual and attention processing (*79–81*). (iv) Sleep Patterns. Eligible participants had regular sleep schedules, with less than two-hour variations in bedtimes and wake-up times between weekdays and weekends, and average sleep durations of six to nine hours. We assessed these patterns using a sleep-wake habits questionnaire (*5, 37, 69, 70*) and the Munich Chronotype Questionnaire (*82*). (v) Health status. We excluded individuals with physical or psychiatric illnesses, those currently on medication, or those suspected of having sleep disorders, based on self-reported questionnaires (*5, 37, 69, 70*). (vi) Age. Participants were required to be between 18-30 years old to control for age-related influences on sleep structure (*83, 84*).

After providing informed consent for participation in this study, all participants completed the Morningness-Eveningness Questionnaire (MEQ)(*85*) and the Pittsburgh Sleep Quality Index Questionnaire (*86*). Results showed that all participants were neither extreme morning nor evening types with MEQ scores ranging from 44 to 69. Additionally, no participants were classified as poor sleepers, as indicated by PSQI scores that did not exceed five points.

### Experimental design

#### Sleep monitoring stage

To ensure consistent sleep quality among participants during the experiment, participants were instructed to adhere to their normal sleep-wake routines, maintaining stable sleep/wake times and durations. We monitored sleep-wake behavior using a sleep log and/or an actigraphy device (GT9X-BT, ActiGraph) for three to seven days before the experiments. If a participant’s sleep-wake schedule varied by more than two hours between workdays and free days during this monitoring period, we rescheduled the sleep experiment. On the day before the experiment, participants were instructed to refrain from alcohol, excessive physical activity, and napping. On the day of the experiment, they were asked to avoid caffeine consumption.

#### Sleep sessions

Sleep sessions occurred in both the adaptation and training and measurement stage. The adaptation stage familiarized participants to the new sleep environment, ensuring undisturbed sleeps in the training and measurement stage. Typically, the adaptation stage is affected by First-Night Effect (FNE), which can disrupt sleep (*32, 33, 35–37, 87, 88*). The training and measurement stage with VPL learning tasks and a sleep session were conducted to see sleep-facilitated effects on VPL.

Participants completed either two (Experiment 1) or three (Experiment 2) sleep sessions in the early afternoon, approximately one week apart, following our established sleep study protocols (*5, 37, 69–71*). This interval was designed so that any effects from a nap in the early afternoon did not carry over and influence the sleep quality in the next session. Each sleep session lasted for 90 minutes. Before each sleep session, we attached polysomnography (PSG) electrodes to the participants. They were then instructed to sleep in an MRI setting while we recorded PSG data.

#### Training and measurement stage

The final session for each participant explored the interaction between sleep and VPL using the TDT (see Texture discrimination task (TDT)). We designed a sequence of sessions to isolate the effects of sleep on VPL as follows: participants engaged in two training sessions, one (Training A) before and one (Training B) after the sleep session, along with four test sessions. Test 1 and Test 2 took place before and after the Training A. Test 3 and Test 4 were conducted before and after the Training B.

In the two training sessions, Training A and Training B, the background orientations were either horizontal or vertical, with each session’s orientation being orthogonal to the other. For example, if Training A had a horizontal (or vertical) background orientation, then Training B had a vertical (or horizontal) background orientation. Previous studies have indicated that these distinct orientations in two TDT training sessions can induce retrograde interference of VPL from Training B to Training A sessions when the two sessions were performed successively or when a nap was inserted between them but did not contain REM sleep (*5, 38, 69*). Therefore, our experiment design expected observable performance changes due to this interference without REM sleep during the nap (*5, 69*).

Test 1 and Test 4 consisted of both Task A and Task B, each with 180 trials. Test 2 and Test 3 included only Task A with 90 trials. Our design aimed to dissect at least two critical elements of sleep-facilitated effects of VPL. First, we evaluated the offline performance gains in Task A by comparing performance between Test 2 and Test 3, without any potential interference from Task B, as participants did not engage with Task B from Training A up to Test 3. Second, we measured the degree of resilience to retrograde interference of Task A learning by examining the performance change between Test 3 and Test 4, considering that Training B could impact the stability of Task A learning.

In addition to the training and test sessions, we conducted short TDT sessions to familiarize and remind participants of the task. Before Training A, participants conducted an introductory session (See Texture discrimination task (TDT)) until consistently achieved a predefined performance level. Additionally, a reminder session (See Texture discrimination task (TDT)) was conducted after a sleep session and before Test 2 to review the TDT task procedure.

The schedule around the sleep session was tightly controlled. Participants completed Test 1 and immediately began the sleep session, during which we collected simultaneous PSG and MRS. After a break of at least 30 minutes (including the time required to remove the EEG cap) to minimize the influence of sleep inertia on task performance, Test 2 was conducted, with a brief two-minute break between each testing and training session.

### Texture discrimination task (TDT)

We used the texture discrimination task (TDT), a standard and widely used VPL task (*72*), for this study. Conducted in a dimly lit room, participants maintained a fixed head position using a chin rest. Visual stimuli were presented on the computer screen 57 cm away. These stimuli were created with MATLAB and Psychtoolbox extensions (*89, 90*).

Each trial began with a 1000 ms fixation point at the center of the screen, followed by a 17 ms target display. After a varied duration of a blank screen, a 100 ms mask display with randomly oriented v-shaped patterns appeared. Participants were instructed to keep their gaze at the center of the screen throughout the stimulus presentation. The target display spanned 19°, featuring a 19 × 19 array of background lines jittered by 0.25°, oriented horizontally or vertically. It included a central letter (‘L’ or ‘T’) for fixation and three diagonal lines (the target array) located peripherally in a trained visual field quadrant (5-9° eccentricity), aligned horizontally or vertically. The trained quadrant, located in the upper right or left visual field, was randomly assigned but remained constant for each participant. Following the mask display, participants reported via keyboard if the central letter was ‘L’ or ‘T’ and whether the orientation of the target array was horizontal or vertical. Auditory feedback indicated the accuracy of the letter task: a 880 Hz beep for correct and a 440 Hz beep for incorrect responses. No feedback was provided for the orientation task.

The time interval between the target and mask onset, known as the stimulus-to-mask onset asynchrony (SOA), was modulated across trials to regulate the task difficulty. Both test and training sessions had six different SOAs ranging from 266 to 33 ms. In a test session, each SOA had 15 trials, resulting in a total of 90 trials per session. The sequence of SOA presentations was pseudo-randomized to minimize learning effects and participants fatigue (*91*). In a training session, there were 60 trials per SOA, totaling 360 trials.

TDT performance was determined by the SOA threshold for 80% correct responses in the orientation task. To compute the SOA threshold, we calculated the correct orientation task response rate for each SOA in a test session. Using the psignifit toolbox (version 4) (*92*), we fitted a cumulative Gaussian psychometric function with a beta-binomial model to account for overdispersion, and estimated the SOA corresponding to 80% accuracy. Incorrect letter task trials were excluded from this SOA threshold calculation.

Changes in TDT performance were determined by the relative shift in SOA threshold values (measured in ms) between test sessions. For example, the improvement over sleep session was quantified as [100 (%) × (SOA_Test1_– SOA_Test2_)/(SOA_Test 1_)].

The introductory session before the first test session comprised three progressively shorter SOA values: 800 ms, 600 ms, and 400 ms. Starting with 800 ms, the session proceeded to 600 ms and 400 ms, each with 20 trials, totaling 60 trials. Each set of 20 trials was equally divided between horizontal and vertical background orientations. The session was repeated until a participant could perform the orientation task with at least 90% accuracy at the 400 ms SOA. The procedure of the reminder session (*5, 69*) was identical to the introductory session but conducted only once and not repeated.

### Sleepiness measurement

Throughout the experiment, particularly during the combined sleep and VPL sessions, we monitored participant sleepiness/alertness using the Stanford Sleepiness Scale (SSS) (*93, 94*) and the Psychomotor Vigilance Task (PVT) (*95*). The SSS ratings ranged from 1 (feeling active, vital, alert or wide awake) to 7 (no longer fighting sleep, sleep onset soon or having dream-like thoughts). Participants selected a rating on the scale that best described their level of sleepiness.

The PVT was administered using a custom MATLAB script with the same parameters as in previous studies (*5, 69*). Each trial began with a fixation screen, followed by a magenta circle at the center of the target screen. Participants were instructed to press the spacebar as quickly as possible upon recognizing the circle. The interval between the fixation and the target screens ranged from 1,000 to 10,000 ms. The PVT duration was approximately 2 min. Reaction times (RTs), expressed in seconds, were log-transformed to reduce the skewness of data distribution. Median RT measured behavioral indicators of sleepiness.

We collected sleepiness data before the four test sessions to ensure that varying sleepiness levels, as indexed by the SSS score and median RTs from the PVT, did not confound our results. We examined differences in these sleepiness measures across the four test sessions in the two groups: DLPFC group and mPFC group. For the SSS, Friedman tests revealed no significant differences in SSS scores across the four time points in either group (DLPFC: *χ*²(3) = 1.88, *p* = 0.60; mPFC: *χ*²(3) = 5.15, *p* = 0.16). For the PVT, Repeated-measures ANOVAs indicated no significant effect of time on PVT performance in either group (DLPFC: *F*(3,42) = 0.64, *p* = 0.59, partial *η*² = 0.04; mPFC: *F*(3,45) = 1.95, *p* = 0.14, partial *η*² = 0.12).

### PSG measurement

PSG consisted of brain waves (Electroencephalography or EEG), eye movements (Electrooculogram or EOG), muscle activity (Electromyogram or EMG), and heart rhythm (Electrocardiogram or ECG). Before recording, it took about 40 minutes to attach the necessary electrodes to each participant. PSG was used to determine the sleep stages (see below) during simultaneous MRS recording. For these measurements, we used the two MRI-compatible PSG caps for the PSG recordings: one with 32 channels and another with 25 channels, provided by Brain Products GmbH.

The 32-electrode cap consisted of 23 EEG electrodes for scalp, six for EOG, two for EMG, and one for ECG electrode. Horizontal EOGs were obtained from two bipolar electrodes at the outer canthi of both eyes, and vertical EOGs from four electrodes above and below the left and right eyes. The 25-electrode configuration included 18 scalp EEG electrodes, four EOG electrodes, two EMG electrodes, and one ECG electrode. For this setup, EOGs were recorded with bipolar electrodes at the outer canthi of each eye, and two additional electrodes above and below the left eye. In both types of caps, EMGs were recorded from the chin in a bipolar manner, and the ECG was obtained from a spot two-finger-widths left of the spine, level with the lower shoulder blades. EEG and ECG electrodes were referenced to Fz, with the ground electrode positioned at AFp4. Electrode impedances were kept at or below 10 kΩ for EEG and ECG, and below 15 kΩ for EOG and EMG. Data acquisitions were performed using an MRI-compatible amplifier (BrainAmp MR, Brain Products) and recording software (BrainVision Recorder, Brain Products) at a sampling rate of 5,000 Hz. Out of the total 31 participants, 26 used the 32-channel EEG caps.

Scanner artifacts and ballistocardiogram artifacts in the PSG data, recorded simultaneously with the MRS, were removed using Brain Vision Analyzer 2 (Brain Products). During denoising, the data were low-pass filtered at 50 Hz and were downsampled to 500 Hz. Subsequently, the EEG data were re-referenced to the left (TP9) and right (TP10) mastoids.

### Sleep-stage scoring and sleep parameters

We scored sleep stages for each 30-second epoch in the American Academy of Sleep Medicine (AASM) guidelines (*96*). The stages included wakefulness (stage W), NREM stage 1 (stage N1), NREM stage 2 (stage N2), NREM stage 3 (stage N3), and REM (stage REM). To describe the sleep architecture in each experiment, we calculated standard sleep parameters (Table 1). These parameters included sleep-onset latency (time to transition into stage N2 after lights off), the percentage of each sleep stage, wake time after sleep onset, sleep efficiency (percentage of time spent asleep relative to the total time in bed), and the total time in bed (interval from lights off to lights on).

**Table 1:**
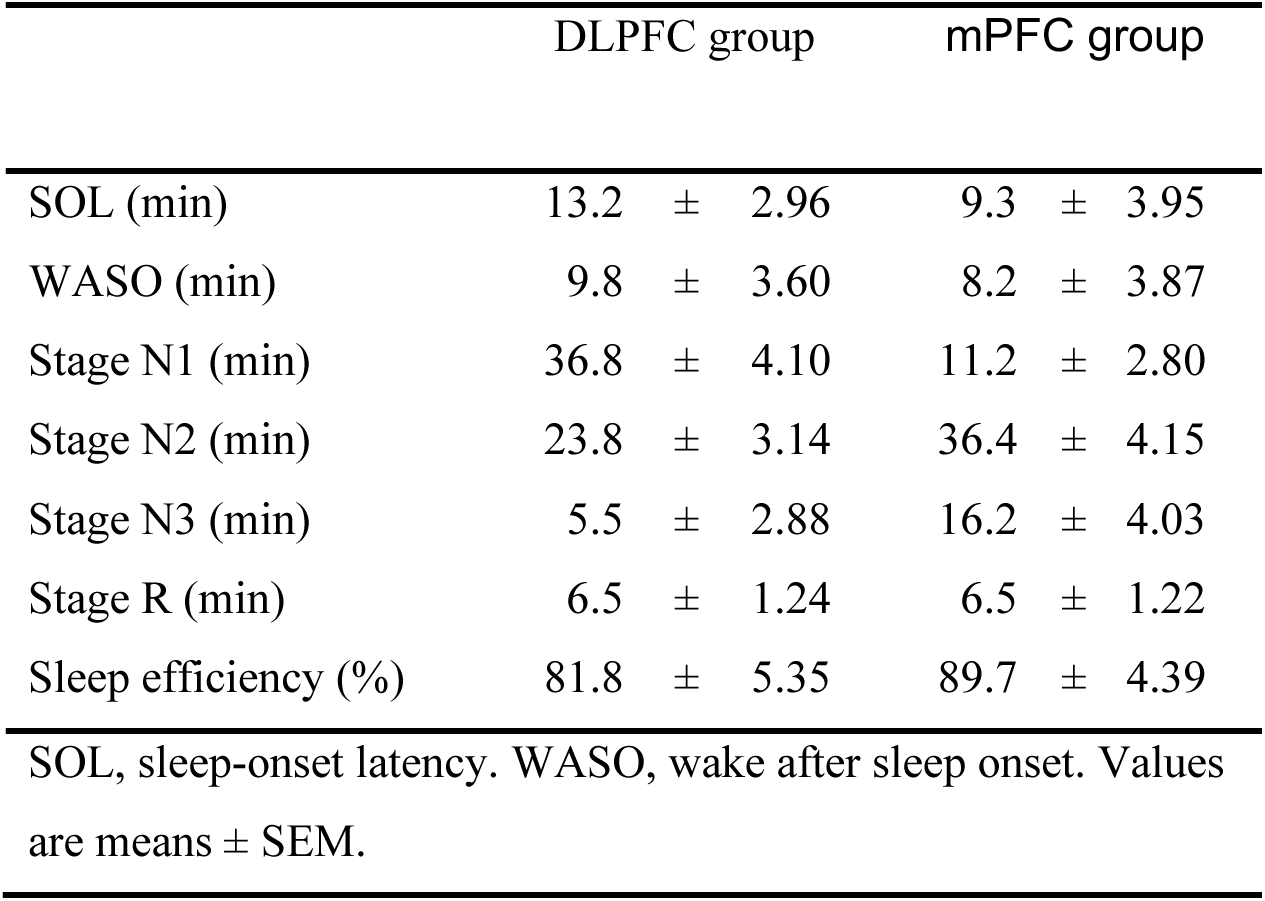
Sleep Structures.

|  | DLPFC group | mPFC group |
| --- | --- | --- |
| SOL (min) | 13.2 ± 2.96 | 9.3 ± 3.95 |
| WASO (min) | 9.8 ± 3.60 | 8.2 ± 3.87 |
| Stage N1 (min) | 36.8 ± 4.10 | 11.2 ± 2.80 |
| Stage N2 (min) | 23.8 ± 3.14 | 36.4 ± 4.15 |
| Stage N3 (min) | 5.5 ± 2.88 | 16.2 ± 4.03 |
| Stage R (min) | 6.5 ± 1.24 | 6.5 ± 1.22 |
| Sleep efficiency (%) | 81.8 ± 5.35 | 89.7 ± 4.39 |
| SOL, sleep-onset latency. WASO, wake after sleep onset. Values are means ± SEM. |  |  |

### MRS acquisition

MRI was performed on a 3T Siemens Prisma scanner using a 64-channel head coil. Participants’ heads were secured with cushions and gauze to prevent motion and discomfort. Additional support, including back and knee cushions, and blankets to maintain body warmth, was provided upon request.

For each sleep session, we first obtained a high-resolution T1-weighted anatomical reference image dataset using a magnetization-prepared rapid gradient echo (MP-RAGE) sequence. This sequence provided detailed brain structure visualization, crucial for subsequent voxel of interest (VOI) placement. The parameters were: time-to-repeat (TR) = 1.9s, time-to-echo (TE) = 3.02 ms, flip angle (FA) = 9°, with a voxel size of 1 × 1 × 1 mm across 256 slices and no slice gap.

We placed VOIs in two prefrontal regions: the medial prefrontal cortex (mPFC) and the dorsolateral prefrontal cortex (DLPFC). For the mPFC, a 2.5 × 2.5 × 2.5 cm³ voxel was positioned bilaterally, immediately anterior to the genu of the corpus callosum, using MP-RAGE images for anatomical guidance. For the DLPFC, we used a slightly smaller 2.2 × 2.2 × 2.2 cm³ voxel on the middle frontal gyrus. The larger voxel in the mPFC than in the DLPFC ensured adequate signal-to-noise ratio (SNR) in this relatively homogeneous gray-matter region, whereas the smaller voxel in the DLPFC reduced contamination from neighboring white matter and sulci, increasing regional specificity. These dimensions match established practice in prefrontal MRS studies (*97–99*). The voxel difference between the mPFC and DLPFC should not be problematic, particularly because the MRS signals during NREM and REM sleep in each region were normalized by those obtained during wakefulness in the same region. We further minimized white matter inclusion to reduce lipid contamination. Shimming, carried out with the scanner’s automated tool, optimized field homogeneity. The entire preparation—structural imaging, VOI placement, and shimming—was completed within 20–30 minutes.

The MEGA-PRESS sequence measured the concentrations of GABA and the combination of glutamate and glutamine (Glx) within the VOI. This technique uses a frequency-selective editing pulse to isolate specific metabolite signals, consistent with established methodologies. The ‘Edit On’ and ‘Edit Off’ phases differentiated GABA and Glx concentrations from the composite spectrum. Two MEGA-PRESS sequences were conducted: an initial short test scan to ensure data quality quickly, followed by a more extensive sleep scan with increased averages for higher accuracy. The test scan parameters were: TR = 1.25 s; TE = 68 ms; FA = 90°; with 32 averages, lasting 2:05 minutes, following the previous MRS-PSG studies (*5, 37, 71*). The sleep scan mirrored these parameters but increased the averages to 240, extending the scan duration to 10:05 minutes. During each sleep session, we performed the longer 10-min scans repeatedly, totaling nine times, to monitor the neurochemical changes across the sleep stages.

### Temporal co-registration of MRS and PSG data

PSG and MRS data obtained during sleep sessions were aligned in time using procedures established in previous studies (*5, 37, 71*). Sleep was defined as three states: wakefulness, NREM sleep (stages N1–N3 combined), and REM sleep. Because PSG epochs were scored every 30 s (*96*), we aggregated them into blocks and assigned each corresponding MRS segment to sleep stage most represented in that block (*5, 37, 71*). This procedure ensured that the neurochemical measures could be interpreted within the context of the prevailing sleep state.

### MRS analysis for E/I ratio during NREM and REM sleep

We measured the levels of neurometabolites in the brain, including Glx, GABA, and NAA. For analysis, we used LCModel (*100, 101*), focusing on a specific chemical shift range of the MRS signal from 1.95 to 4.2 ppm. We used NAA as a control metabolite to calculate Glx and GABA concentrations, following procedures from earlier research (*5, 25*). The LCModel operates under the assumption that the obtained spectrum can be fitted to a linear combination of individual metabolite spectra from an imported basis set (*100, 101*).

After the co-registration of MRS data with the PSG data in time, we quantified the levels of E/I ratios, as the concentration of Glx divided by that of GABA. This analysis involved two primary steps: First, we calculated the average levels of E/I ratio during wakefulness, NREM and REM sleep, respectively. The average levels of E/I ratio during wakefulness – defined as the period from lights off to sleep onset and any instances of wake after sleep onset – worked as the baseline. Second, we calculated the changes in these E/I ratio levels during NREM and REM sleep relative to wakefulness baseline. For example, the E/I ratio during NREM sleep was computed using the formula: [(E/I_non-REM_ – E/I_wakefulness_)/E/I_wakefulness_ x 100(%)]. This formula provides the percentage change of E/I ratio during NREM and REM sleep relative to the wakefulness baseline.

### Quality tests for the MRS data

We assessed MRS data quality using four parameters established in prior studies (*5, 37*): Cramer–Rao lower bounds (CRLB), shim values, NAA linewidth, and frequency drift. CRLB, which measure fitting error, served as the main exclusion criterion for low-quality spectra, with a threshold of 25% (*26, 98, 102*). Shim values (<30 Hz considered optimal (*103*)) reflected magnetic field homogeneity. Both DLPFC and mPFC groups showed mean shim values well below this threshold (5.0 ± 0.24 Hz and 7.3 ± 0.43 Hz, respectively). NAA linewidths—a measure of spectral resolution—averaged 9.2 ± 0.16 Hz (DLPFC) and 10.5 ± 0.45 Hz (mPFC). Volunteer MRS studies typically report NAA linewidths of 9–11 Hz, values considered acceptable and indicative of high spectral quality (*99*). Frequency drift, reflecting B0 instability and potential head motion, remained low in both groups (2.7 ± 1.02 Hz, DLPFC; 2.3 ± 0.32 Hz, mPFC) and within reported acceptable ranges (*97, 104*). As shown in Table 2, all four indices fell within appropriate limits in both regions, consistent with published findings.

**Table 2.**
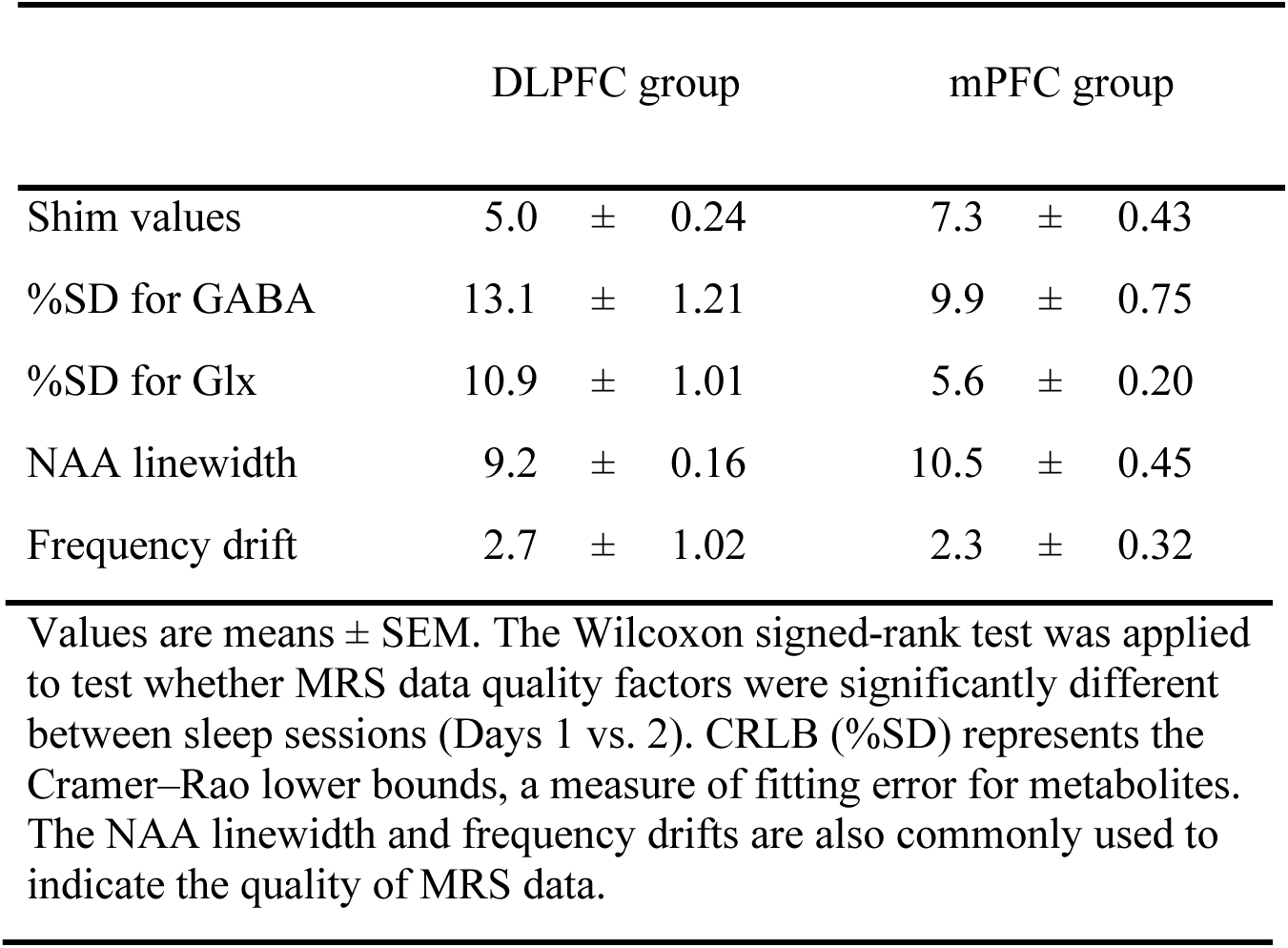
MRS data quality.

| DLPFC group | mPFC group |
| --- | --- |

|  |  |  |  |  |  |  |
| --- | --- | --- | --- | --- | --- | --- |
| Shim values | 5.0 | ± | 0.24 | 7.3 | ± | 0.43 |
| %SD for GABA | 13.1 | ± | 1.21 | 9.9 | ± | 0.75 |
| %SD for Glx | 10.9 | ± | 1.01 | 5.6 | ± | 0.20 |
| NAA linewidth | 9.2 | ± | 0.16 | 10.5 | ± | 0.45 |
| Frequency drift | 2.7 | ± | 1.02 | 2.3 | ± | 0.32 |

Values are means ± SEM. The Wilcoxon signed-rank test was applied to test whether MRS data quality factors were significantly different between sleep sessions (Days 1 vs. 2). CRLB (%SD) represents the Cramer–Rao lower bounds, a measure of fitting error for metabolites. The NAA linewidth and frequency drifts are also commonly used to indicate the quality of MRS data.

### Statistical analysis

We used two-tailed tests (α=0.05) and report mean ± s.e.m. with percentile-bootstrap 95% CIs. Because several variables failed Shapiro–Wilk tests, we analyzed all primary contrasts with resampling. One-sample and paired effects were tested with bootstrap t-tests (2,000 resamples; subject-level case resampling or resampling of paired differences). Correlations and repeated-measures ANOVAs used 2,000 bootstrap resamples to form empirical sampling distributions; two-sided P values equal the fraction of replicates as or more extreme than the observed statistic. Behavioral measures—offline performance gains and resilience to retrograde interference from TDT SOA thresholds—were tested against zero separately in the DLPFC (Experiment 1) and mPFC (Experiment 2) cohorts. For MRS, we defined E/I change as the percent shift from wakefulness to NREM or REM and tested each stage against zero, then related E/I changes to behavior with Pearson correlations whose CIs and P values came from bootstrap resampling. We quantified source-level gamma connectivity (LORETA) between EVA and prefrontal sites and entered values into a 2 × 2 repeated-measures ANOVA (Pathway: DLPFC–EVA vs mPFC–EVA; Sleep Stage: NREM vs REM) with bootstrap inference; significant interactions prompted planned paired post hoc bootstrap t-tests (NREM vs REM within pathway). We checked alertness with Friedman tests on SSS scores and repeated-measures ANOVAs on PVT median log-RTs (Greenhouse–Geisser when needed). Spectra with CRLB>25% were excluded; Day-1 vs Day-2 shim, NAA linewidth, and frequency drift were compared by Wilcoxon signed-rank tests. We identified outliers with median-based modified z-scores and removed them only from the implicated analysis (counts detailed in Results/legends); REM-dependent analyses included only participants with REM-labeled MRS/PSG segments. Effect sizes are Cohen’s d, partial η², Pearson’s r, and matched rank-biserial correlation; we did not correct for multiplicity because tests addressed a priori hypotheses. TDT thresholds were fit with psignifit v4 (beta-binomial); statistics were run in SPSS v27, R v4.3.0, GraphPad Prism v10.6.0, and JASP 0.95.3.

## Author contributions

TY and YS designed the study. TY and TLC performed the experiments. TY analyzed the data. TY, TW, and YS wrote the manuscript.

## Conflict of Interests

The authors declare no competing financial interests.

## Data availability

The datasets generated and/or analyzed in the current study will be available as source data. The computer code that was used to generate results central to the conclusions of this study is available from the corresponding author upon request.

## Acknowledgments

This work was supported by the NIH (R01EY031705, R01EY019466, R01EY027841), NSF (NSF-BSF: 2241417) and KAKENHI (JP20KK0268).

## Supplementary information

**Supplementary Figure 1.**
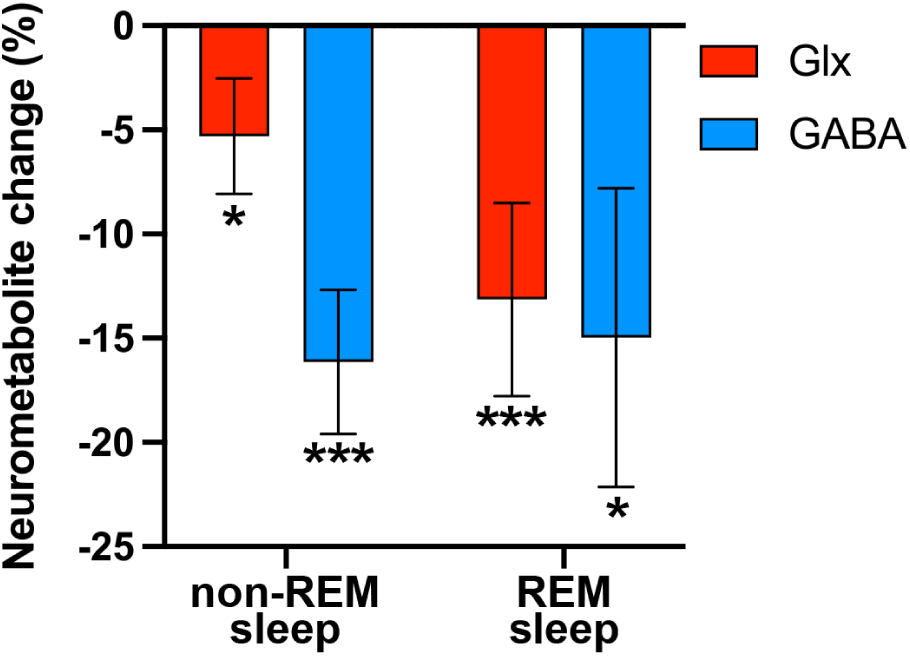
Mean (±s.e.m.) Glx and GABA changes from wakefulness to NREM sleep (left) and REM sleep (right) in DLPFC. The Shapiro–Wilk tests showed that the Glx data for NREM and REM sleep met the normality assumption (NREM sleep: W=0.92, *p*=0.23; REM sleep: *W*=0.88, *p*=0.15). The GABA data for NREM and REM sleep met the normality assumption (NREM sleep: *W*=0.93, *p*=0.26; REM sleep: *W*=0.92, *p*=0.38). To unify the method, we conducted the bootstrap analysis. Based on 2,000 resamples, the results yielded the following results: Glx decreased significantly from wakefulness to both NREM *(p*=0.029, Cohen’s *d*=-0.53, 95% CI=[-10.73, -0.47]) and REM sleep (*p*<0.001, d = -0.95, 95% CI = [-22.12, -5.11]). GABA also decreased significantly from wakefulness to NREM (p<0.001, Cohen’s *d*=-1.25, 95% CI = [-22.34, -9.60]) and REM sleep (*p*=0.014, Cohen’s *d*=-0.70, 95% CI = [-28.26, -1.95]). \**p*<0.05, \*\**p*<0.01, \*\*\**p*<0.001.

**Supplementary Figure 2.**
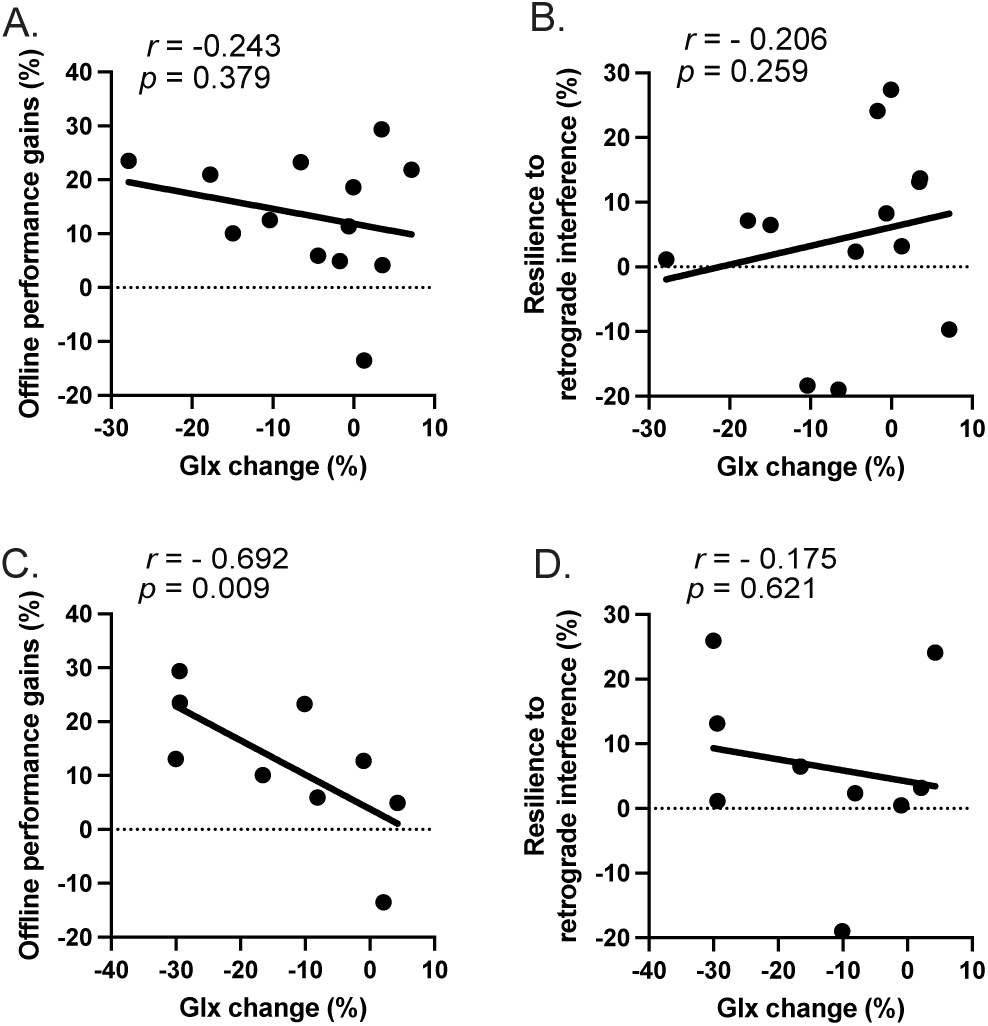
Scatter plots in Experiment 1. A. Offline performance gains vs. changes in Glx level in DLPFC from wakefulness to NREM sleep (*r*=–0.243, *t*=-0.829, *p*=0.379, 95% CI=[-0.697, 0.307]). B. Resilience to retrograde interference gains vs. changes in Glx level in DLPFC from wakefulness to NREM sleep (*r*=-0.206, *t*=0.697, *p*=0.259, 95% CI=[-0.203, 0.591]). C. Offline performance gains vs. changes in Glx level in DLPFC from wakefulness to REM sleep (*r*=-0.692, *t*=-2.536, *p*=0.009, 95% CI=[-0.916, -0.315]). D. Resilience to retrograde interference gains vs. changes in Glx level in DLPFC from wakefulness to REM sleep (*r*=-0.175, *t*=-0.472, *p*=0.621, 95% CI=[-0.845, 0.554]). Solid lines represent linear regression fits obtained using the least-squares method. Correlation coefficients, p-values, and 95% confidence intervals were estimated using bootstrap resampling with 2,000 iterations.

**Supplementary Figure 3.**
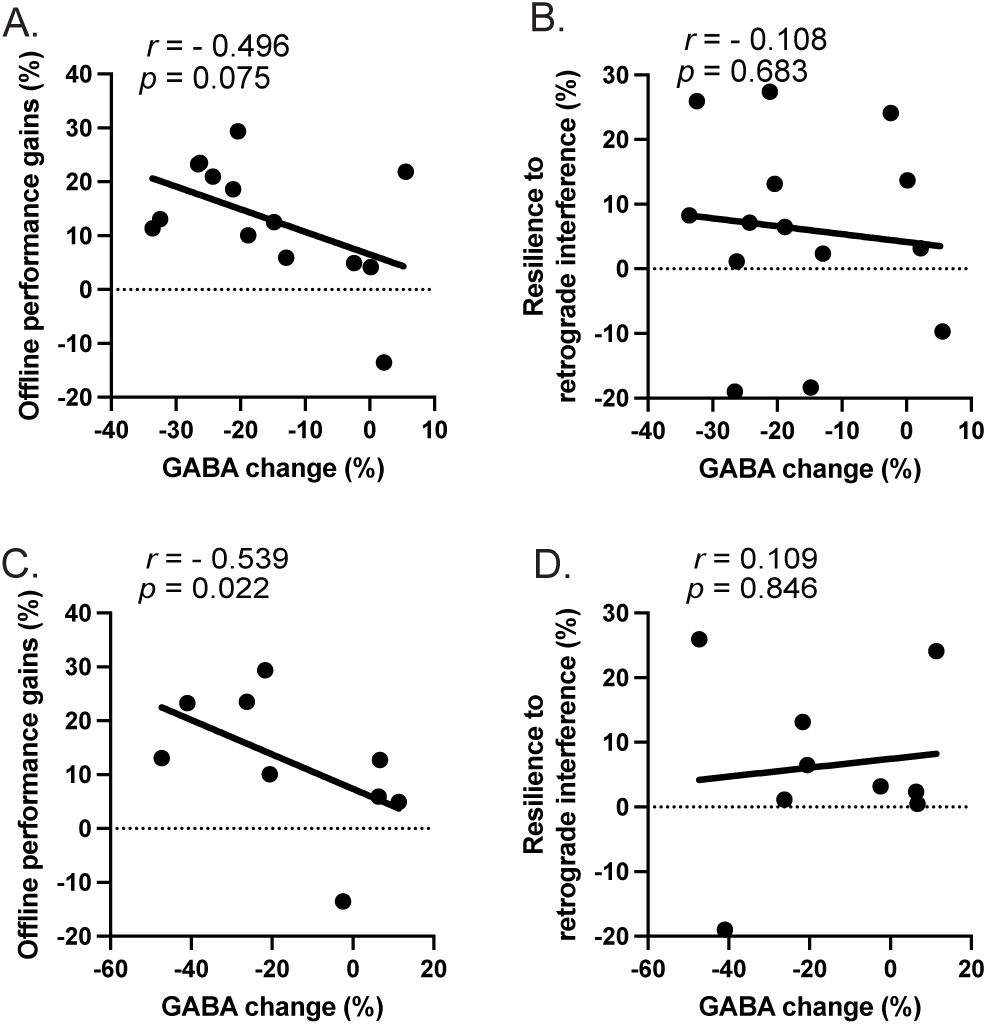
Scatter plots in Experiment 1. A. Offline performance gains vs. changes in GABA level in DLPFC from wakefulness to NREM sleep (*r*=–0.496, *t*=-1.977, *p*=0.075, 95% CI=[-0.882, 0.086]). B. Resilience to retrograde interference gains vs. changes in GABA level in DLPFC from wakefulness to NREM sleep (*r*=-0.108, *t*=-0.375, *p*=0.683, 95% CI=[-0.582, 0.415]). C. Offline performance gains vs. changes in GABA level in DLPFC from wakefulness to REM sleep (*r*=-0.539, *t*=-1.691, *p*=0.022, 95% CI=[-0.875, -0.227]). D. Resilience to retrograde interference gains vs. changes in GABA level in DLPFC from wakefulness to REM sleep (*r*=0.109, *t*=0.289, *p*=0.846, 95% CI=[-0.910, 0.864]). Solid lines represent linear regression fits obtained using the least-squares method. Correlation coefficients, p-values, and 95% confidence intervals were estimated using bootstrap resampling with 2,000 iterations.

**Supplementary Figure 4.**
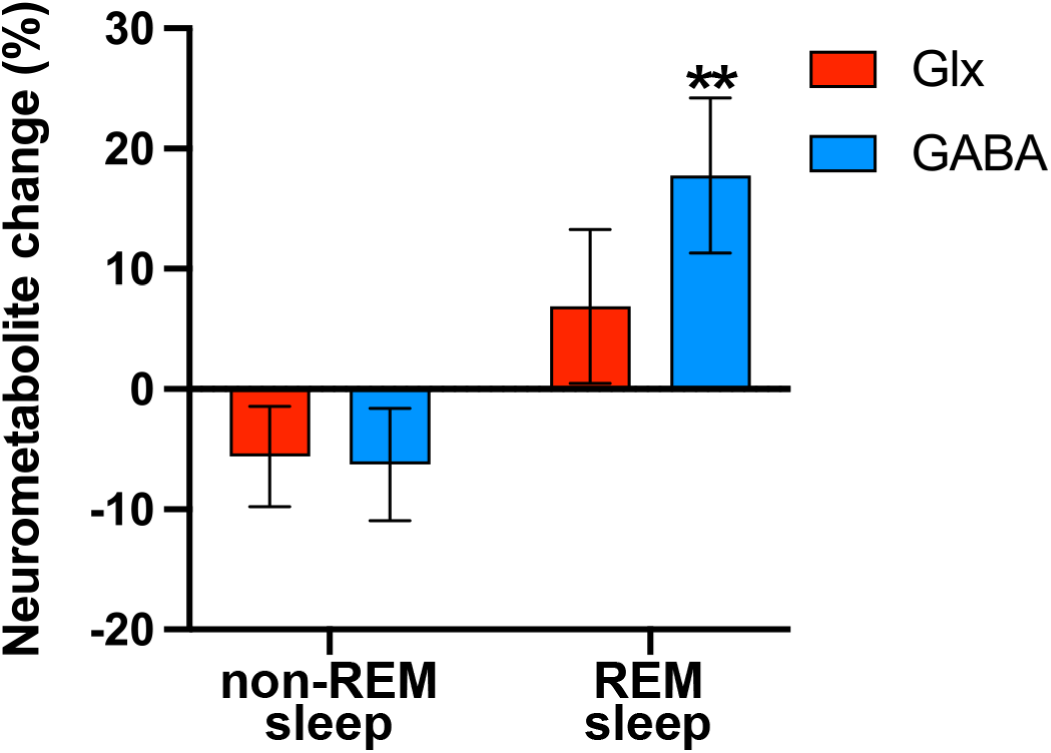
Mean (±s.e.m.) Glx and GABA changes from wakefulness to NREM sleep (left) and REM sleep (right) in mPFC. The Shapiro–Wilk tests showed that the Glx data for NREM and REM sleep met the normality assumption (NREM sleep: *W=*0.95, *p*=0.55; REM sleep: *W*=0.95, *p*=0.62). The GABA data for NREM and REM sleep met the normality assumption (NREM sleep: *W*=0.94, *p*=0.29; REM sleep: *W*=0.94, *p*=0.54). To unify the method, we conducted the bootstrap analysis. Based on 2,000 resamples, the results yielded the following results: Glx changes were not significant in either NREM (*p=0*.17, Cohen’s *d*=-0.34, 95% CI=[-13.54, 2.32]) or REM sleep (*p*=0.26, Cohen’s *d*=0.31, 95% CI = [-5.19, 18.56]). GABA changes were not significant in NREM (*p*=0.19, Cohen’s *d*=-0.33, 95% CI = [-14.52, 2.62]) but were significant in REM sleep (*p*=0.004, Cohen’s *d*=0.79, 95% CI=[5.96, 29.20]). \*\**p*<0.01.

**Supplementary Figure 5.**
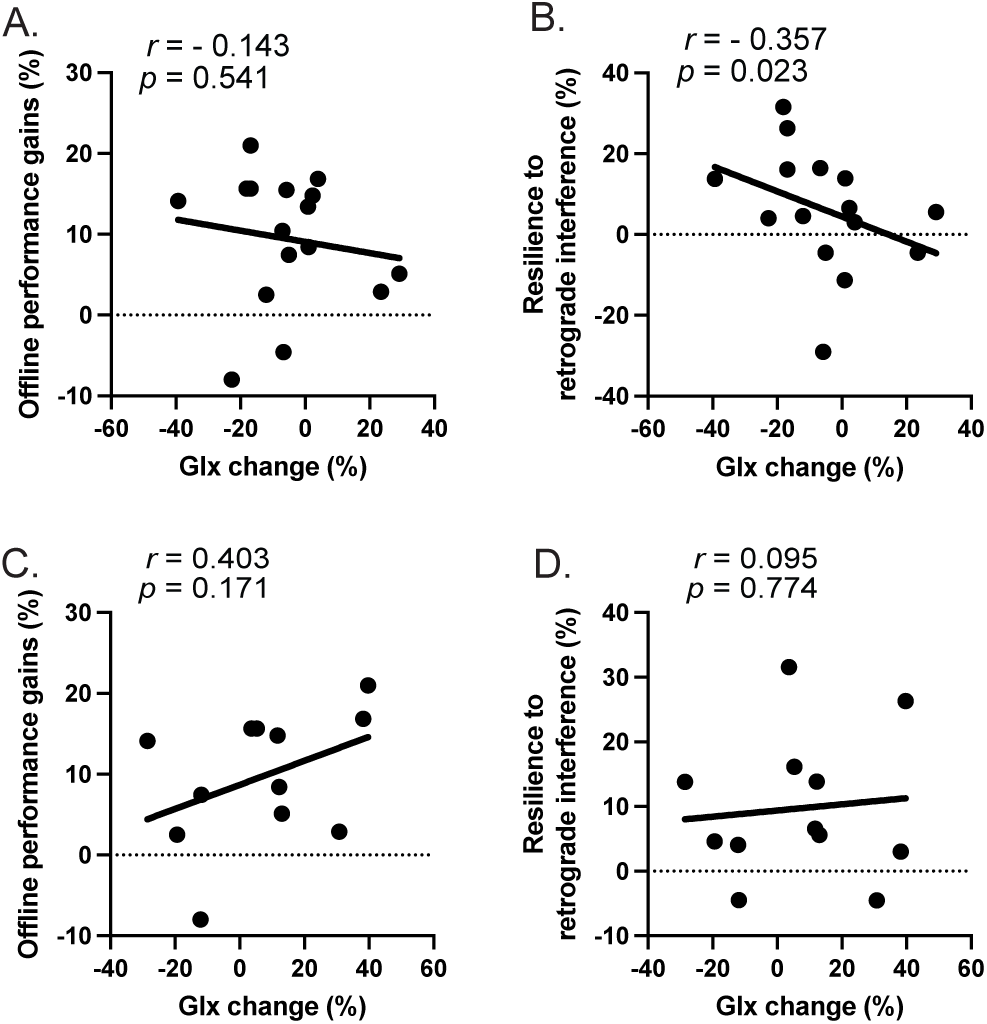
Scatter plots in Experiment 2. A. Offline performance gains vs. changes in Glx level in DLPFC from wakefulness to NREM sleep (*r*=–0.143, *t*=-0.539, *p*=0.541, 95% CI=[-0.678, 0.364]). B. Resilience to retrograde interference gains vs. changes in Glx level in DLPFC from wakefulness to NREM sleep (*r*=-0.357, *t*=-1.379, *p*=0.023, 95% CI=[-0.673, -0.062]). C. Offline performance gains vs. changes in Glx level in DLPFC from wakefulness to REM sleep (*r*=0.403, *t*=1.394, *p*=0.171, 95% CI=[-0.245, 0.800]). D. Resilience to retrograde interference gains vs. changes in Glx level in DLPFC from wakefulness to REM sleep (*r*=0.095, *t*=0.302, *p*=0.774, 95% CI=[-0.531, 0.666]). Solid lines represent linear regression fits obtained using the least-squares method. Correlation coefficients, p-values, and 95% confidence intervals were estimated using bootstrap resampling with 2,000 iterations.

**Supplementary Figure 6.**
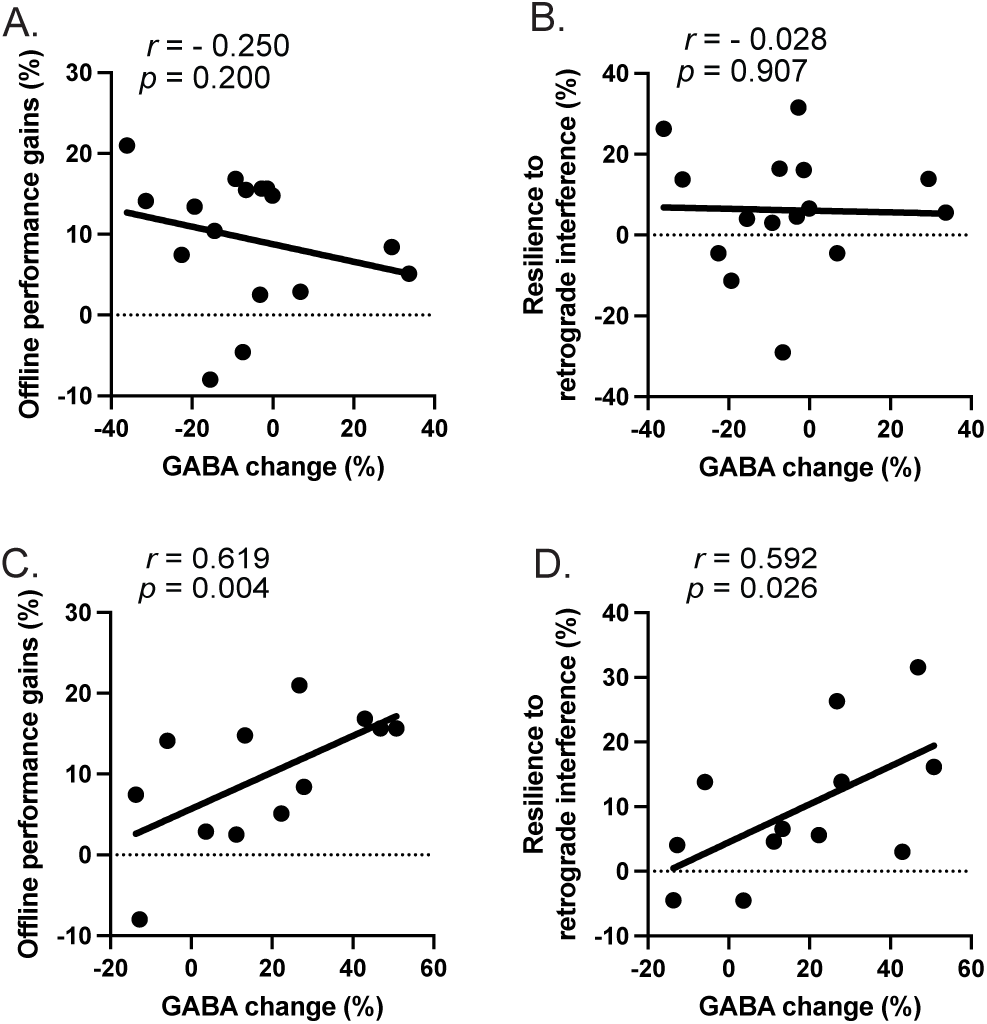
Scatter plots in Experiment 2. A. Offline performance gains vs. changes in GABA level in DLPFC from wakefulness to NREM sleep (*r*=–0.250, *t*=-0.968, *p*=0.200, 95% CI=[-0.657, 0.130]). B. Resilience to retrograde interference gains vs. changes in GABA level in DLPFC from wakefulness to NREM sleep (*r*=-0.028, *t*=-0.101, *p*=0.907, 95% CI=[-0.447, 0.413]). C. Offline performance gains vs. changes in GABA level in DLPFC from wakefulness to REM sleep (*r*=0.619, *t*=2.493, *p*=0.004, 95% CI=[0.227, 0.888]). D. Resilience to retrograde interference gains vs. changes in GABA level in DLPFC from wakefulness to REM sleep (*r*=0.592, *t*=2.323, *p*=0.026, 95% CI=[0.099, 0.877]). Solid lines represent linear regression fits obtained using the least-squares method. Correlation coefficients, p-values, and 95% confidence intervals were estimated using bootstrap resampling with 2,000 iterations.

## Notes

### Competing Interest Statement

The authors have declared no competing interest.

### Summary of Updates

We removed a descriptive paragraph explaining the rationale for the additional adaptation session in Experiment 2, as it was not essential for understanding the experimental design or interpretation of the results. This change does not affect the data, analyses, or conclusions of the study.

